# *FAS^lpr^* gene dosage tunes the extent of lymphoproliferation and T cell differentiation in lupus

**DOI:** 10.1101/2023.11.01.563607

**Authors:** Ritu Bohat, Xiaofang Liang, Yanping Chen, Chunyu Xu, Ningbo Zheng, Ashley Guerrero, Roshni Jaffery, Nicholas A. Egan, Adolfo Robles, M. John Hicks, Yong Du, Xiqun Chen, Chandra Mohan, Weiyi Peng

## Abstract

*Sle1* and *Fas^lpr^* are two lupus susceptibility loci that lead to manifestations of systemic lupus erythematosus. To evaluate dosage effects of *FAS^lpr^* in determining cellular and serological phenotypes associated with lupus, we developed a new C57BL/6 (B6) congenic lupus strain, B6.*Sle1/Sle1.Fas^lpr^*^/+^ (sle1^homo^.lpr^het^) and compared it with B6.*Fas^lpr/lpr^*(lpr^homo^), B6.*Sle1/Sle1* (sle1^homo^), and B6.*Sle1/Sle1.Fas^lpr/lpr^* (sle1^homo^.lpr^homo^) strains. Whereas Sle1^homo^.lpr^homo^ mice exhibited profound lymphoproliferation and early mortality, sle1^homo^.lpr^het^ mice had a lifespan comparable to B6 mice, with no evidence of splenomegaly or lymphadenopathy. Compared to B6 monogenic lupus strains, sle1^homo^.lpr^het^ mice exhibited significantly elevated serum anti-dsDNA antibodies and increased proteinuria. Additionally, Sle1^homo^.lpr^het^ T cells had an increased propensity to differentiate into Th1 cells. Gene dose effects of *Fas^lpr^* were noted in upregulating serum IL-1α, IL-2, and IL-27. Taken together, sle1^homo^.lpr^het^ mice emerge as a more faithful model of human SLE, ideal for genetic studies, autoantibody repertoire investigation, and for exploring Th1 effector cell skewing.

## 1. Introduction

Systemic lupus erythematosus (SLE) is a chronic, prototypic autoimmune disorder which could result in damage to a wide range of organs, including skin, kidneys, and the neural system [1]. Besides heterogeneous clinical manifestations, the high complexity of SLE is also reflected by a variety of risk factors. Among them, multiple genetic factors have been associated with the predisposition to SLE [2]. Earlier studies revealed that certain MHC alleles are consistent high-risk factors in SLE patients, which could be related to MHC restriction for T cell autoantigens [3]. Further studies to link single-gene mutations with SLE susceptibility have identified serval high-risk polymorphisms, including those that cause dysregulations in DNA clearance, type I interferon (IFN), or lymphocyte activation pathways [2]. It pinpoints that self-antigen generation and hyper-activation of innate and adaptive immunity are the major players in SLE pathogenesis.

In addition to clinical studies, congenic murine models bearing lupus susceptibility loci have been widely employed to dissect the contributions of different genetic aberrations to manifestations of SLE [4]. *Sle1* and *Fas^lpr^* are two lupus susceptibility loci associated with the well-researched spontaneous lupus models, NZM2410 and MRL/*lpr,* respectively [5]. Subsequently, these two loci were reported to result in dysfunctions of the adaptive immunity and/or *FAS/FASL*-mediated apoptosis pathway, respectively [6, 7]. However, individually introducing (the NZM2410 allele of) *Sle1* and *Fas^lpr^* onto a lupus-resistant C57BL/6 (B6) background only led to mild autoimmune phenotypes. For example, renal pathological changes are largely negligible in B6*.Fas^lpr/lpr^* (lpr^homo^) mice before 9-month age [8, 9]. Previous studies further demonstrated the epistatic interaction of *Sle1* and *Fas^lpr^* results in severe lupus in the B6 background. High levels of anti-nuclear and anti-glomerular autoantibodies can be detected in 3-month B6.*Sle1.Fas^lpr/lpr^* (sle1^homo^.lpr^homo^) mice. This strain exhibits massive lymphosplenomegaly and glomerulonephritis as early as 2-3 months of age, suggesting that *Sle1* and *Fas^lpr^*impact non-redundant pathways leading to lymphoproliferative autoimmunity [10]. Although the sle1^homo^.lpr^homo^ strain could well mirror human SLE by driving lymphoproliferative autoimmunity; most SLE patients exhibit mild or no clinical evidence of lymphoproliferation [11]. Given the dominant impact of the *FAS/FASL* axis on lymphocyte apoptosis [12, 13], we investigated whether *Fas^lpr^* gene dosage on the sle1^homo^ background might play a role in the extent of lymphoproliferation in this lupus model.

To address this, we comprehensively characterized lymphoproliferation status and other cellular and serological phenotypes associated with lupus in a new congenic strain, B6.*Sle1/Sle1.Fas^lpr^*^/+^ (sle1^homo^.lpr^het^). Importantly, our studies suggest that this novel strain with a reduced dosage of *Fas^lpr^* is a more faithful model of human SLE based on its serological and renal phenotypes without the confounding effect of lymphoproliferation. Whereas the complete absence of *FAS* leads to impaired activation-induced cell death, reduced levels of *FAS* regulate T cell differentiation and production of key cytokines implicated in autoimmunity, particularly Th1-related cytokines.

## 2. Materials and methods

### 2.1 Mice

C57BL/6 (B6) mice and B6 mice homozygous *FAS^lpr/lpr^* were obtained from The Jackson Laboratory (#000482, Bar Harbor, ME). B6 congenic strains bearing the NZM2410-derived *sle1* gene interval and *FAS^lpr^*were generated as previously described [7] and subsequently bred in a specific pathogen-free barrier facility at the University of Houston. Mice were handled following protocols approved by Institutional Animal Care and Use Committees. Equal numbers of male and female mice were used, and any observed sex differences were indicated.

### 2.2 Serum collection and analyses

100µl of serum samples were collected from experimental mice at different time points and stored at -80°C for batch analyses to determine anti-dsDNA (Anti-double-stranded DNA) autoantibody and cytokine/chemokine levels. The anti-dsDNA ELISA was performed as previously described [7]. Briefly, serum samples were collected from experimental mice and stored at -80°C for future batch sample processing. To measure serum levels of cytokine/chemokines in congenic lupus strains, a Luminex assay using the ProcartaPlex™ Mouse Cytokine & Chemokine Panel 1 36-plex kit (#EPXR360-26092-901, ThermoFisher Scientific, Waltham, MA) was performed according to manufacturer’s instructions. A Luminex 200 Xmap analyzer (Luminex Corporation, Austin, TX) was used to determine the serum levels of tested soluble factors based on the magnitude of fluorescent signal in a microparticle-specific manner. Blood Urea Nitrogen (BUN) was also determined in the same serum samples using a urea colorimetric assay kit (#MAK006, Sigma-Aldrich, St. Louis, MO).

### 2.3 Urine sample collection and analyses

Experimental mice were individually housed in mouse metabolic cages (#MM030505-10R, Lenderking caging products, Millersville, MD) for 24 hours to collect urine samples. The 24-hour (hr) urine volume was recorded, and urine samples were stored at -80°C and analyzed to determine urinary protein concentration by a Bradford assay using the Pierce detergent compatible kit (#23246, ThermoFisher Scientific). The 24-hr urine protein amount was calculated based on 24-hr urine volume and total urinary protein concentration.

### 2.4 Splenocytes collection and flow cytometry analyses

Single-cell suspensions of spleen tissues were prepared and depleted of red blood cells by ACK lysing buffer (#A1049201, ThermoFisher Scientific). For flow cytometric analyses, splenocytes were washed twice with staining buffer and incubated with a cocktail of antibodies targeting surface markers at 4°C for 30 min. Cells were then fixed and permeabilized using either the Foxp3 /transcription factor staining buffer set (#00-5523-00, ThermoFisher Scientific) or the BD fixation/permeabilization solution kit (#554714, BD BioSciences, Franklin Lakes, NJ) according to the manufacturer’s protocols and then incubated with a cocktail of antibodies against intracellular markers. The antibodies used for staining included anti-CD4-PB (Thermo Fisher Scientific Cat# 48-0042-82, RRID:AB_1272194), anti-CD8-PE/cy7 (Tonbo Biosciences Cat# 60-0081 (also OWL-A07793), RRID:AB_2621832), anti-CD25-Percp (Tonbo Biosciences Cat# 65-0251 (also OWL-A11282), RRID:AB_2621889), anti-FoxP3-PE (Thermo Fisher Scientific Cat# 12-5773-82, RRID:AB_465936), anti-CD19-APC (BD Biosciences Cat# 550992, RRID:AB_398483); (Tonbo Biosciences Cat# 65-0193 (also OWL-A11277), RRID:AB_2621887), anti-Ki67-FITC (Thermo Fisher Scientific Cat# 11-5698-82, RRID:AB_11151330); (Thermo Fisher Scientific Cat# 25-5698-82, RRID:AB_11220070), anti-CD62L-PE (Cytek Biosciences Cat# 65-0621, RRID:AB_3073622), anti-CD44-PB (Tonbo Biosciences Cat# 35-0441 (also OWL-A05062), RRID:AB_2621688), anti-NK1.1-APC (BD Biosciences Cat# 550627, RRID:AB_398463), (Thermo Fisher Scientific Cat# 48-0112-82, RRID:AB_1582236), anti-CD11c-APC (Tonbo Biosciences Cat# 20-0114 (also OWL-A00976), RRID:AB_2621557), anti-F4/80-PE (Tonbo Biosciences Cat# 50-4801 (also OWL-A05860), RRID:AB_2621795), anti-Ly6G-PE/cy7 (Tonbo Biosciences Cat# 60-1276 (also OWL-A07821), RRID:AB_2621860), anti-Ly6C-Percp (Thermo Fisher Scientific Cat# 45-5932-82, RRID:AB_2723343), anti-CD19-percp (#65-0193, TONBO Biosciences) anti-CD5-PE (Cytek Biosciences Cat# 50-0051, RRID:AB_3073623), anti-CD21-APC (BioLegend Cat# 123412 (also 123411), RRID:AB_2085160), anti-CD23-PB (BioLegend Cat# 101616 (also 101615), RRID:AB_2103306), anti-Ki67-PE/Cy7 (Thermo Fisher Scientific Cat# 25-5698-82, RRID:AB_11220070), anti-CD93-PE (BioLegend Cat# 136503 (also 136504), RRID:AB_1967094), and anti-IgM-FITC (BioLegend Cat# 406505 (also 406506), RRID:AB_315055). An BD LSRFortessa X-20 cell analyzer (RRID:SCR_019600) was used for acquisition.

### 2.5 T cells differentiation

Naïve CD4^+^ T cells were isolated from single-cell suspensions from spleen tissue by negative selection using the EasySep Mouse CD4^+^ T cell Isolation Kit (# 19852, StemCell Technologies, Cambridge, MA) and differentiated into different T helper (Th) subsets. Purified naïve CD4^+^ T cells were stimulated with 2.5 μg/ml of anti-CD3e (Tonbo Biosciences Cat# 70-0031 (also OWL-A11492), RRID:AB_2621472) and 1 μg/ml of anti-CD28 (BD Biosciences Cat# 553294, RRID:AB_394763) antibodies in different types of culture medium for T cell differentiation and proliferation [14-16]: (a) Th0 condition: MEM*a* medium (#32-571-101, Fisher scientific, Hampton, NH) supplemented with 10% FBS (#S11150, R&D Systems, Minneapolis, MN), 100U/ml of IL-2 (NDC 65483-116-07, Prometheus laboratories, San Diego, CA, USA); (b) Th1 condition: MEM*a* medium supplemented with 10% FBS, 100U/ml of IL-2, 4 ng/ml of IL-12 (# 210-12, PEPRO TECH, East Windsor, NJ), and 10 μg/ml of anti-IL-4 mAb (Bio X Cell Cat# BE0045 RRID:AB_1107707); (c) Th2 condition: MEM*a* medium supplemented with 10% FBS, 100U/ml of IL-2, 10 ng/ml of IL-4 (#214-14, PeproTech), and 10 μg/ml of anti-IFN-γ mAb (Bio X Cell Cat# BE0055 RRID:AB_1107694); (d) Th17 condition: MEM*a* medium supplemented with 10% FBS, 100U/ml of IL-2, 30ng/ml of IL-6 (#216-16, PeproTech), 20ng/ml of IL-1p (#211-11B PeproTech), 10 μg/ml of anti-IFN-γ mAb, 2.5 ng/ml of TGF-ý1 (#763104, BioLegend).

To evaluate T cell proliferation, stimulated T cells were cultured in MEM*a* medium supplemented with 10% FBS and 100U/ml of IL-2. To determine the percentage of differentiated Th1, Th2, and Th17, differentiated T cells were restimulated with 50ng/ml of PMA (#P1585, Sigma-Aldrich) and 1μg/ml of ionomycin (#I0634, Sigma-Aldrich) for 1 hour, with blockade of cytokine secretion being achieved by simultaneous treatment with 4 µl of GolgiStop (# 51-2092KZ, BD Biosciences) for every 6 ml of cell culture for the last 3 hours. Restimulated T cells were stained with panels of antibodies for flow cytometric analysis as described previously. Additional antibodies used for this experiment included anti-IFNγ-APC (Tonbo Biosciences Cat# 20-7311 (also OWL-A01038), RRID:AB_2621616), anti-IFNγ-PE (Tonbo Biosciences Cat# 50-7311 (also OWL-A05875), RRID:AB_2621810), anti-IL-4-PE (BioLegend Cat# 504102 (also 504101), RRID:AB_315316), anti-IL-4-PB (Thermo Fisher Scientific Cat# 48-0042-82, RRID:AB_1272194), and anti-IL-17A-FITC (Thermo Fisher Scientific Cat# 11-7177-81, RRID:AB_763581). To measure cytokine production by differentiated T cells, T cells were re-seeded in a new 96-well plate and incubated overnight with fresh culture medium in the presence of 50 ng/ml of PMA. The concentration of IFN-γ, IL-4, and IL-17 in the culture supernatant was determined using ELISA kits from R&D systems DY485, DY404, and DY421, respectively.

### 2.6 Histopathology

Mice were sacrificed at 4-8 months of age to collect kidneys. Kidney samples were Formalin-Fixed and Paraffin-Embedded (FFPE). FFPE tissue slides were prepared and stained with Hematoxylin and Eosin (H&E). Sections were analyzed using a light microscope and scored for evidence of glomerulonephritis, and tubulo-interstitial disease, as detailed previously [17]. Kidneys were critically analyzed for any signs of cell proliferation and infiltration, thickening, and fibrous changes.

### 2.7 Statistical analyses

Summary statistics (e.g., mean, SEM) of the data are reported. Assessments of differences in continuous measurements between the two groups were made using the two-sample t-test. Either one-way or two ANOVA was performed to determine the significance in various subsets. The Kaplan-Meier method and log-rank test were used to compare survival between groups. p-value of less than 0.05 was considered significant. Data acquisition and analysis from the cell analyzer were executed on BD FACSDiva Software (RRID:SCR_001456) and BD FlowJo software (RRID:SCR_008520). Graph generation and statistical analyses were performed using GraphPad Prism software (RRID:SCR_002798).

## 3. Results

### 3.1 Sle1^homo^.lpr^homo^ mice, but not Sle1^homo^.lpr^het^ mice, exhibited lymphosplenomegaly with reduced survival

To further characterize the interaction between the two lupus-associated loci, *Sle1* and *Fas^lpr^,* on regulating autoimmune responses in the C57BL/6 (B6) background, B6.*Sle1/Sle1* (sle1^homo^) mice were crossed with B6.*Fas^lpr/lpr^*(lpr^homo^) mice over 2 generations and selected for B6.*Sle1/Sle1.Fas^lpr^*^/+^ (sle1^homo^.lpr^het^) and B6.*Sle1/Sle1.Fas^lpr/lpr^*(sle1^homo^.lpr^homo^) mice. As illustrated in Figures 1A and 1B, weights and cell numbers of spleens and inguinal nodes in sle1^homo^.lpr^homo^ mice at 2-3 months (mo) of age were significantly higher than those in B6, and other tested B6 congenic mice. We observed 100% penetrance of enlarged lymph nodes in sle1^homo^.lpr^homo^ mice during their life span (Fig 1C). All tested sle1^homo^.lpr^homo^ mice achieved primary experimental endpoint before or at 10-mo of age, marked primarily by severe lymphadenopathy (long diameter of any lymph node >15mm; Fig 1D). In contrast, enlarged spleen and lymph nodes were not noted in sle1^homo^.lpr^het^, lpr^homo^ and sle1^homo^ mice. The survival curves from these two B6 congenic strains were comparable to the wild-type B6 strain. These results indicate that the complete loss of *FAS* function is required to elicit lymphosplenomegaly and accelerated mortality in B6 mice bearing the *sle1* locus.

**Figure 1.**
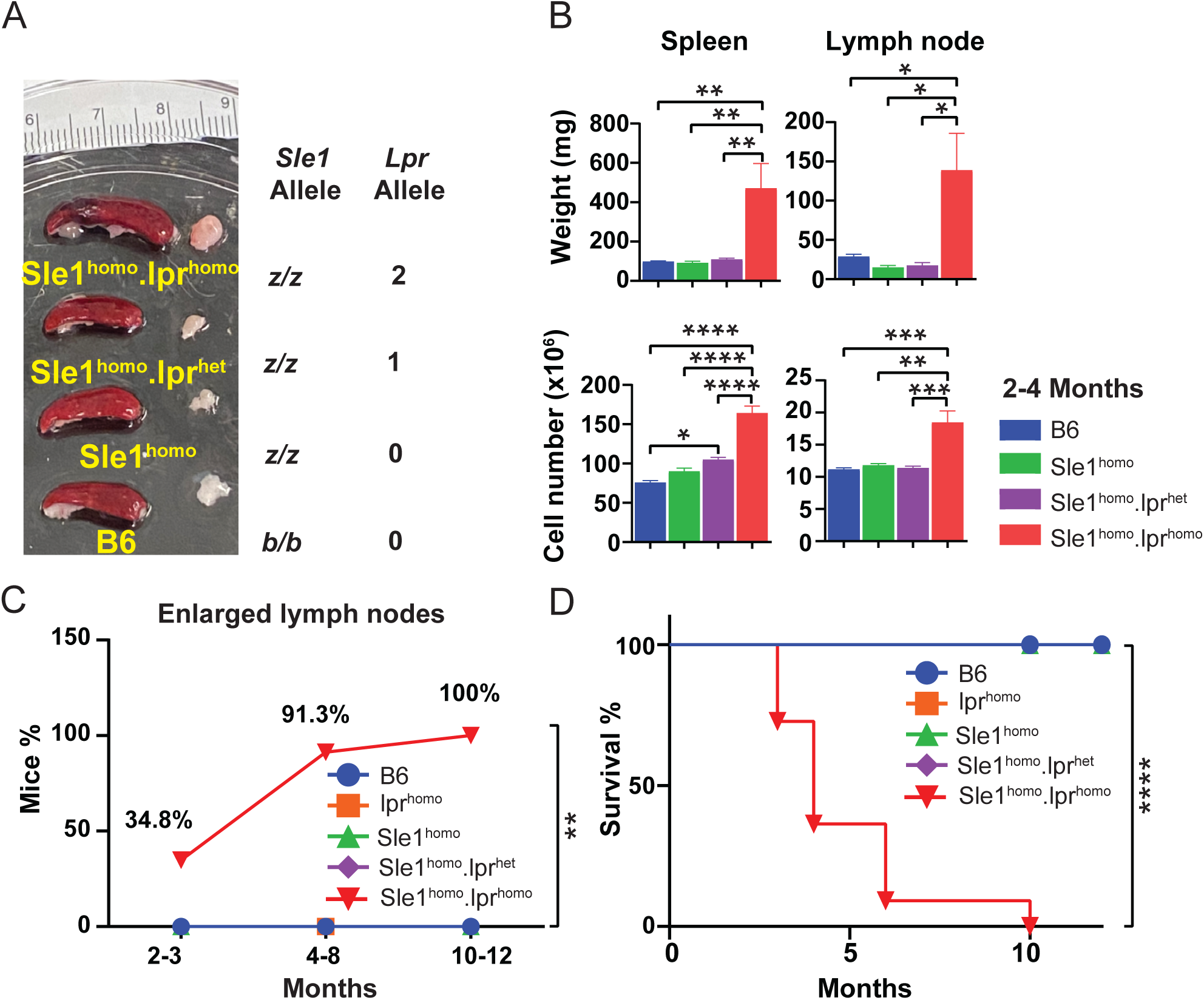
Sle1^homo^.lpr^homo^, but not Sle1^homo^.lpr^het^ mice exhibit prominent splenomegaly, lymphadenopathy, and reduced survival. (A) Representative image of spleens and inguinal lymph nodes from B6 mice and B6 congenic mice at the age of 3 months (mo). The B6 and NZM240 alleles of *Sle1* are indicated as the *b* and *z* alleles, respectively. The numbers of *Lpr* alleles in *Fas* loci are also listed for each B6 congenic strain. (B) Weight and total cell numbers in the spleen and an inguinal lymph node. (C) Lymphadenopathy percentages: The lymph node size and health conditions of experimental mice were monitored at least twice per week. Mice with visible lymph nodes were counted as having lymphadenopathy. The endpoints include any lymph node >1.5 cm or severe health issues. (D) Survival data. Tested B6 congenic mice include Sle1^homo^, lpr^homo^, Sle1^homo^.lpr^het^, and Sle1^homo^.lpr^homo^. N=4-6/strain; Mice from both genders were included. *p<0.05, **p<0.01, ***p<0.001 and ****p<0.0001.

### 3.2 Acceleration of autoimmune responses by loss of *FAS* function in B6.*sle1* congenic mice is gene-dose dependent

Next, we examined whether complete loss of *FAS* function is required to promote autoimmune responses in B6 mice bearing the *sle1* locus homozygously. B6 and B6 congenic mice were examined for serum levels of anti-nuclear antibody (ANA), urine protein levels, and histopathological changes in renal tissues at three different ages: 2-3 mo, 4-8 mo, and 10-12 mo. Given that the mortality in sle1^homo^.lpr^homo^ mice older than 4 months is high, and this strain has been well characterized previously [10], we only included 2-3 mo-old sle1^homo^.lpr^homo^ mice in comparative studies. Compared with age-matched B6 controls, 2-3 mo-old sle1^homo^.lpr^homo^ mice exhibited significantly elevated serum anti-dsDNA IgG levels, while marginal increases were found in sle1^homo^ and sle1^homo^.lpr^het^ mice (Fig 2A; Supplementary Tab 1). Moreover, serum ANA levels in sle1^homo^.lpr^het^ mice at the age of 4-8 mo were significantly higher than the ANA levels in sle1^homo^ and B6 controls (Fig 2A; Supplementary Tab 1). We also started to observe elevated ANA levels in sle1^homo^ mice at the age of 10-12 mo. Increased proteinuria was observed in sle1^homo^.lpr^homo^, sle1^homo^.lpr^het^ and sle1^homo^ mice starting at the age of 2-3 mo, 4-8 mo and 10-12 mo, respectively (Fig 2B; Supplementary Tab 2). Higher levels of BUN were found in all B6 congenic strains than the B6 controls in all three age groups, with the phenotype being more prominent in sle1^homo^.lpr^homo^ (Fig 2C; Supplementary Tab 3). Consistent with the marginal increase in proteinuria and BUN in sle1^homo^.lpr^het^ mice compared to the sle1^homo^ controls, both these groups of mice demonstrated mild glomerulonephritis (GN) at the age of 4-8mo, as illustrated in part in Table 1 and Figure 2D. These alterations include foci of inflammatory cell infiltrates and fibrous tissue formation in glomerular and peri-glomerular regions. This contrasts with the severe (Class IV) GN noted in sle1^homo^.lpr^homo^ mice reported previously [10]. Our results suggest that homozygosity in both *Sle1* and *Fas^lpr^* is required to cause severe renal tissue damage. On the other hand, partial loss of *FAS* function promotes autoimmune responses and accelerates the earlier emergence of serum ANA and proteinuria in B6 mice congenic for the *sle1* locus.

**Figure 2.**
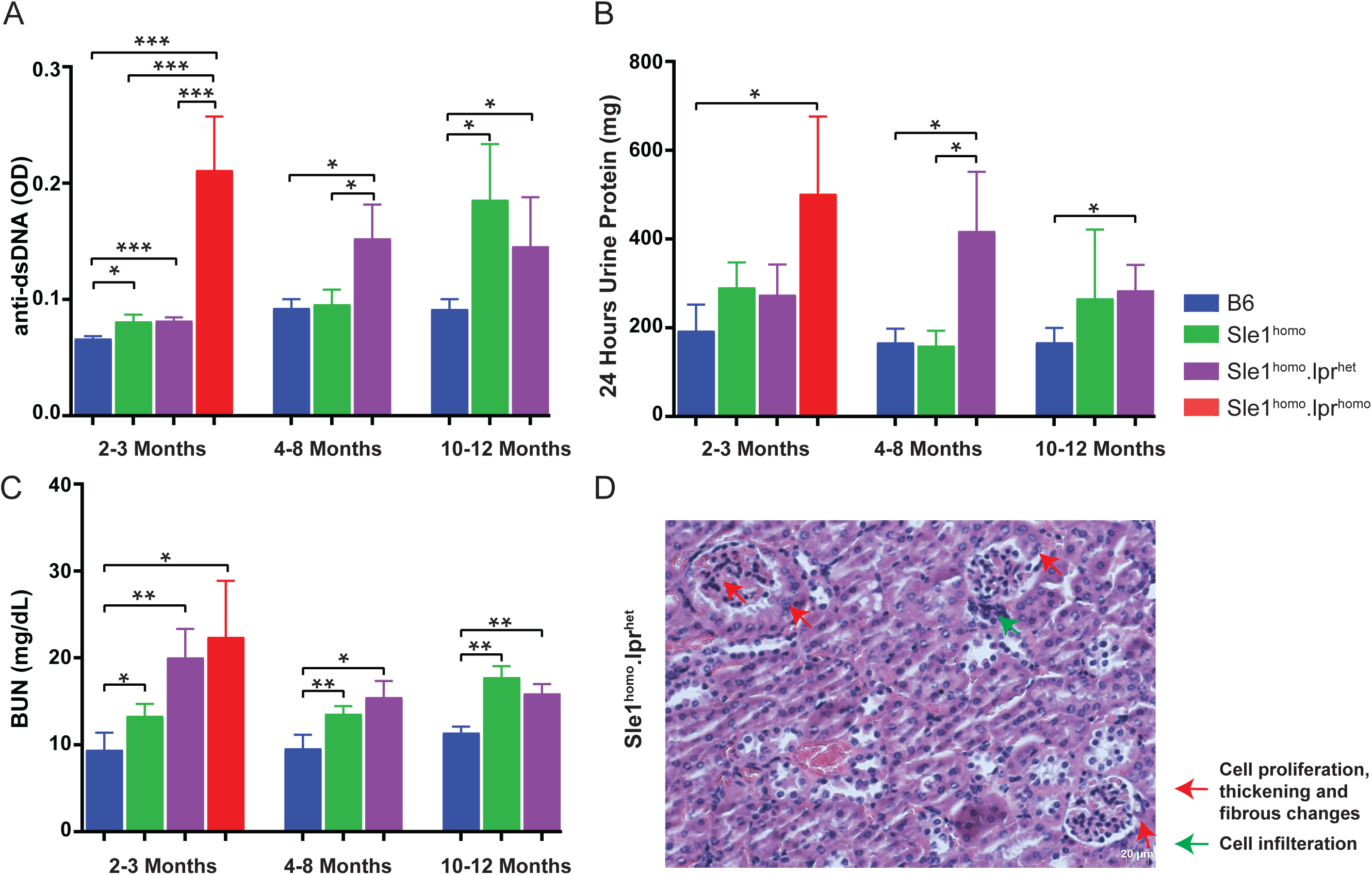
Impact of *FAS^lpr^* zygosity on the levels of serum autoantibody, blood urea nitrogen, and urine protein. (A) Serum levels of IgG autoantibodies to dsDNA in B6 and B6 congenic mice. Sera were collected from experimental mice at three different ages: 2-3 months (mo), 4-8 mo, and 10-12 mo, and were examined for anti-dsDNA IgG by ELISA. (B) Total urine protein amount. The 24-hr urine volume and protein concentration were measured and used to calculate the total amount of 24-hr urine protein. (C) Blood Urea Nitrogen (BUN) levels in B6 and B6 congenic mice. Tested B6 congenic mice include Sle1^homo^, Sle1^homo^.lpr^het^, and Sle1^homo^.lpr^homo^. Mice from both genders were included. *p<0.05, **p<0.01, ***p<0.001 and ****p<0.0001. (D) Typical renal changes observed in Sle1^homo^.lpr^het^ mice. Kidney tissues from 4-8 mo old B6 congenic mice were stained with H&E and examined for evidence of any pathological changes. A representative image of renal tissues from Sle1^homo^.lpr^het^ mice is shown.

**Table 1.** Summary of histopathology data of kidney samples. B6 and B6 congenic mice were sacrificed at 4-8 mo of age to collect kidneys. Scores for evidence of glomerulonephritis and tubulo-interstitial disease in FFPE kidney tissue slides are shown.

| Mice strain | Gender | Age | Severity of GN<br>Grade (0-4) | Glomeruli with Crescent<br>formation (%) | Tubulo-interstitial<br>nephritis pathological<br>damage<br>Grade (0–4) | GlomerGlomeruli (any sclerosis,<br>collapse or thrombonecrotic<br>lesions) Grade (0-4) |
| --- | --- | --- | --- | --- | --- | --- |
| B6-1 | F | 8 Months | 0 | 0 | 0 | 0 |
| B6-2 | F | 8 Months | 0 | 0 | 0 | 0 |
| B6-3 | F | 8 Months | 0 | 0 | 0 | 0 |
| Lpr<sup>homo</sup> -1 | F | 4 months | 0 | 0 | 0 | 0 |
| Lpr<sup>homo</sup> -2 | F | 4 months | 0 | 0 | 0 | 0 |
| Lpr<sup>homo</sup> -3 | F | 4 months | 0 | 0 | 0 | 0 |
| Sle1<sup>homo</sup> -1 | F | 6 months | 1 | 0 | 0 | 0 |
| Sle1<sup>homo</sup> -2 | F | 6 months | 1 | 0 | 0 | 0 |
| Sle1<sup>homo</sup> -3 | F | 6 months | 1 | 0 | 0 | 0 |
| Sle1<sup>homo</sup>.Lpr<sup>het</sup> -1 | F | 8 Months | 1 | 0 | 0 | 0 |
| Sle1<sup>homo</sup>.Lpr<sup>het</sup> -2 | F | 8 Months | 2 | 0 | 0 | 0 |
| Sle1<sup>homo</sup>.Lpr<sup>het</sup> -3 | F | 8 Months | 1 | 0 | 0 | 0 |

**Table 2.**
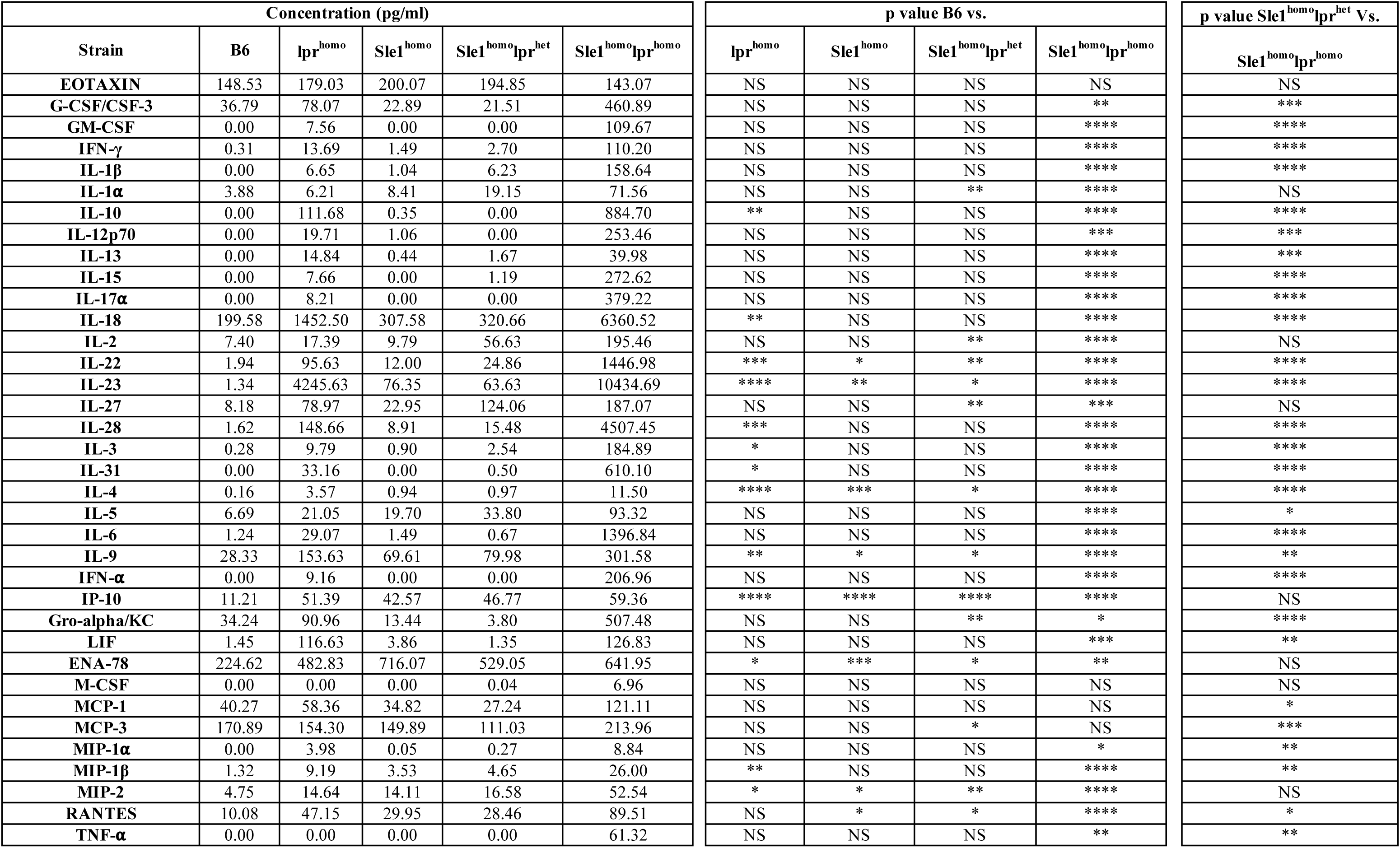
Serum levels of 36 cytokines/chemokines in B6 and B6 congenic mice at the age of 3 months. Averages of serum levels of cytokines/chemokines in each strain and the status of statistical significance between B6 and B6 congenic mice were listed. NS: no statistical significance. *p<0.05, **p<0.01, ***p<0.001 and ****p<0.0001.

### 3.3 Profound alterations in the lymphocyte compartment of peripheral lymphoid organs observed in Sle1^homo^.lpr^homo^ mice, but not Sle1^homo^.lpr^het^ mice

To characterize the immune profiles of peripheral lymphoid organs in B6 and B6 congenic strains, spleen tissues and inguinal lymph nodes from 3-mo-old female and male mice for each strain were collected and examined. As lpr^homo^ mice have been well characterized [8, 18] and have no sign of glomerulonephritis at the age of 4 months (Tab 1), we focused on B6 congenic strains with homozygous *sle1* locus for immune profiling. Phenotypic analysis of splenocytes revealed significant alterations in the cellular composition of major immune cells only in sle1^homo^.lpr^homo^ mice, including reduced percentages of CD8^+^ T cells, B cells, and NK cells, when compared to the B6 controls (Fig 3A and 3B; Supplementary Fig 2). Whereas the major changes in absolute numbers in spleens from sle1^homo^.lpr^homo^ mice included significantly increased CD4^+^ T cells, and numbers of CD8^+^ T cells and B cells in sle1^homo^.lpr^homo^ mice also exhibited a tendency to increase (Supplementary Tab 4).

**Figure 3.**
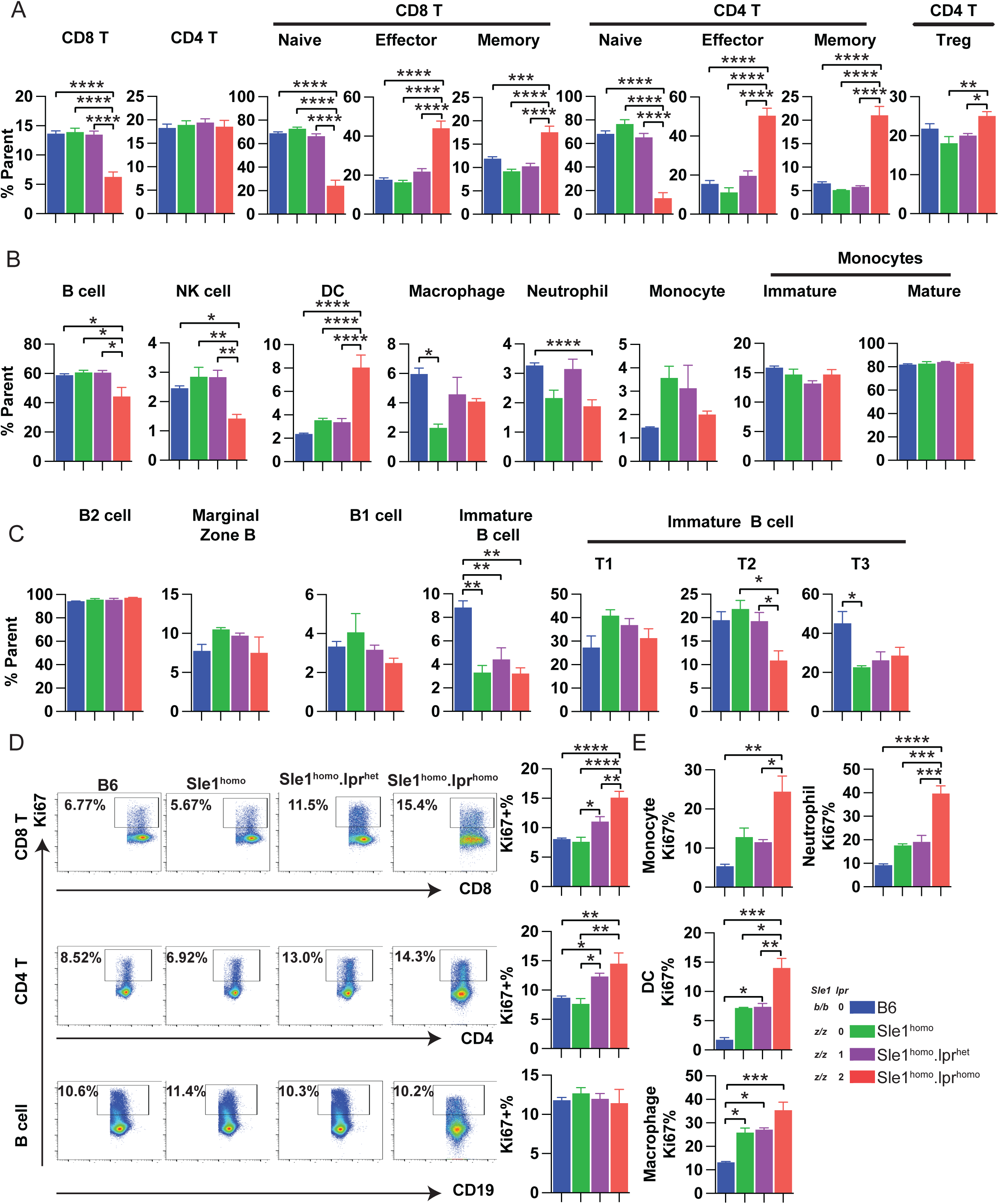
Immune profiling of splenocytes from Sle1^homo^.lpr^homo^ and Sle1^homo^.lpr^het^ mice. Splenocytes were isolated from spleens of B6 and B6 congenic mice at 3 mo of age and stained with a panel of fluorescence-conjugated antibodies. Flow cytometry was used to characterize the abundance and proliferation of immune cells. (A) abundance of T cell subsets, (B) abundance of non-T immune subsets, (C) abundance of B cell subsets, (D) proliferation of lymphocytes, and (E) proliferation of non-T/B immune subsets among different strains are depicted. Tested B6 congenic mice include Sle1^homo^, Sle1^homo^.lpr^het^, and Sle1^homo^.lpr^homo^. B6 mice were used as a control. Mice from both genders were included. N=4-5/strain. *p<0.05, **p<0.01, ***p<0.001 and ****p<0. 0001.

Furthermore, we examined subsets within splenic CD8^+^ T cell, CD4^+^ T cell, and B cell populations. Both CD8^+^ T cells and CD4^+^ T cells in sle1^homo^.lpr^homo^ spleens were skewed towards effector and memory subsets with three-fold lower naïve T cell percentages compared to the B6 controls (Fig 3A). In addition, sle1^homo^.lpr^homo^ mice exhibited marginally increased Treg percentages in spleen tissues (Fig 3A). There were no significant changes in T cell subsets among the remaining B6 congenic strains and B6 controls. Interestingly, sle1^homo^.lpr^homo^ mice exhibited a dramatic increase in splenic cells within the lymphocyte gate that were negative for CD4^+^, CD8^+^, B cell, and NK cell markers (Supplementary Fig 2). Extrapolating from the published literature [10], these are likely to represent CD4^-^ CD8^-^ double negative T cells, which is also consistent with the reduced percentages of CD8^+^ T cells in this strain (Fig 3A).

Unlike T cells, alterations in the B cell compartment in sle1^homo^.lpr^homo^ mice were limited to slightly reduced B1 and immature B cells (Fig 3C). In contrast, the composition of B cell subsets in sle1^homo^.lpr^het^ was largely comparable with B6 controls and sle1^homo^ mice (Fig 3C). The changes in the absolute numbers of immune cells in sle1^homo^.lpr^het^ mice were minor or not statistically significant (Supplementary Tab 4). Moreover, we determined the proliferation status of different immune cells based on the expression of Ki67, a proliferation marker. Both sle1^homo^.lpr^homo^ and sle1^homo^.lpr^het^ spleens revealed increased percentages of proliferative CD8^+^ and CD4^+^ T cells, monocytes, and neutrophils, with more prominent phenotypes being observed in sle1^homo^.lpr^homo^ spleens (Fig 3D and 3E). However, limited or no changes in percentages of proliferative B cells, dendritic cells, and macrophages were seen in all other B6 congenic strains when compared with B6 controls (Fig 3E). Furthermore, homozygosity in both *Sle1* and *Fas^lpr^* resulted in similar changes in the T cell, B cell, and non-lymphoid cell compartments in the lymph nodes, just as those observed in the spleen, while the sle1^homo^.lpr^het^ mice consistently displayed mild phenotypic changes in all immune cells in the lymph nodes (Fig 4A-4C; Supplementary Fig 3; Supplementary Tab 5). Interestingly, unlike moderately increased T cell proliferation observed in spleen tissues, the proliferation of CD8^+^ and CD4^+^ T cells in lymph nodes from sle1^homo^.lpr^het^ remain unchanged compared to the B6 controls (Fig 4D). In contrast, the T and B cells in lymph nodes from sle1^homo^.lpr^homo^ mice displayed significantly increased proliferative status (Fig 4D and 4E).

**Figure 4.**
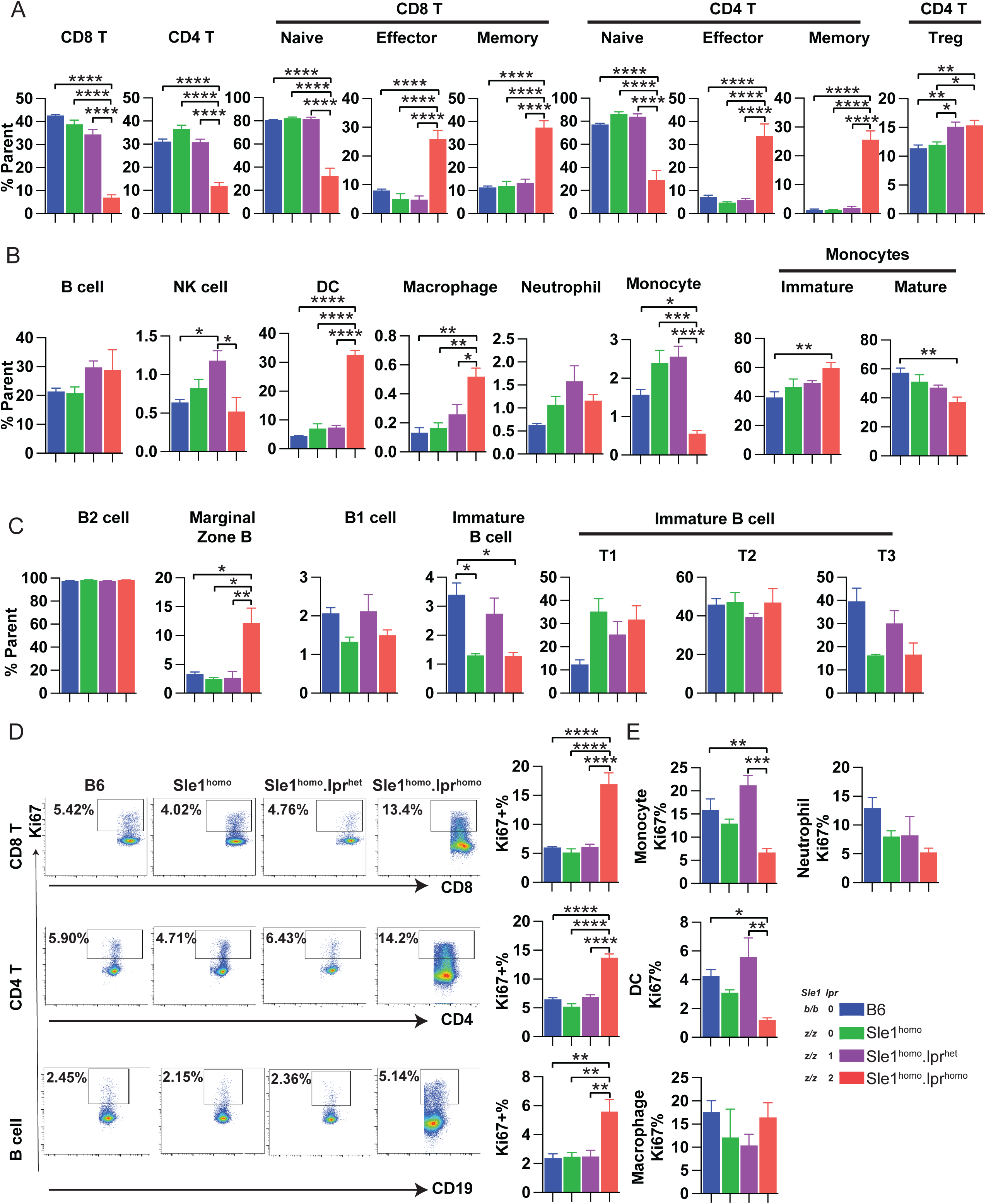
Immune profiling of lymph nodes from Sle1^homo^.lpr^homo^ and Sle1^homo^.lpr^het^ mice. Single-cell suspension samples were prepared from inguinal lymph nodes of B6 and B6 congenic mice at 3 mo of age and stained with a panel of fluorescence-conjugated antibodies. Flow cytometry was used to characterize the abundance and proliferation of immune cells. (A) abundance of T cell subsets, (B) abundance of non-T immune subsets, (C) abundance of B cell subsets, (D) proliferation of lymphocytes, and (E) proliferation of non-T/B immune subsets among different strains are depicted. Tested B6 congenic mice include Sle1^homo^, Sle1^homo^.lpr^het^, and Sle1^homo^.lpr^homo^. N=4-5/strain; C57/B6 (B6) mice were used as control. *p<0.05, **p<0.01, ***p<0.001 and ****p<0.0001.

### 3.4 Gene dose effects of *Fas^lpr^* in regulating serum cytokine levels

Given that several cytokines, such as IL-6, IL-10, IFN-γ and TNF-α, have been implicated in SLE pathogenesis and associated with SLE clinical activity and serological changes [19], we next determined the serum levels of 36 cytokines in B6 and B6 congenic mice at the age of 3 months. Among the tested cytokines, serum levels of 32 cytokines were significantly higher in sle1^homo^.lpr^homo^ mice than those in B6 controls (Tab 2). Unlike sle1^homo^.lpr^homo^ mice, sle1^homo^.lpr^het^ mice exhibited significant elevation in serum levels of fewer cytokines, including IL-1α, IL-2, IL-22, IL-23, IL-27, IL-4, IL-9, IP-10, KC, ENA-78, MCP-3, MIP-2 and RANTES relative to the B6 controls (Tab 2). While the majority of these serum elevations in sle1^homo^.lpr^het^ are shared by sle1^homo^ and/or lpr^homo^ mice, increased IL-1α, IL-2, and IL-27 are only observed in B6 mice with both *Sle1* and *Fas^lpr^*loci, with homozygosity at *Sle1* and *Fas^lpr^* resulting in more profound elevations (Fig 5A). Furthermore, we also observed another group of cytokines whose levels were primarily affected by homozygosity at *Fas^lpr^*. Specifically, lpr^homo^ and sle1^homo^.lpr^homo^ mice, but not sle1^homo^ and sle1^homo^.lpr^het^ mice showed significantly increased IL-3, IL-10, IL-18, IL-28, and IL-31(Fig 5 B and Tab 2). Homozygosity at *Sle1* results in more profound elevations in this group of cytokines. The interaction effect of *Fas^lpr^* and *Sle1* in serum cytokine profiles further supports the notion that *Fas^lpr^* and *Sle1* have non-redundant roles in promoting autoimmune responses.

**Figure 5.**
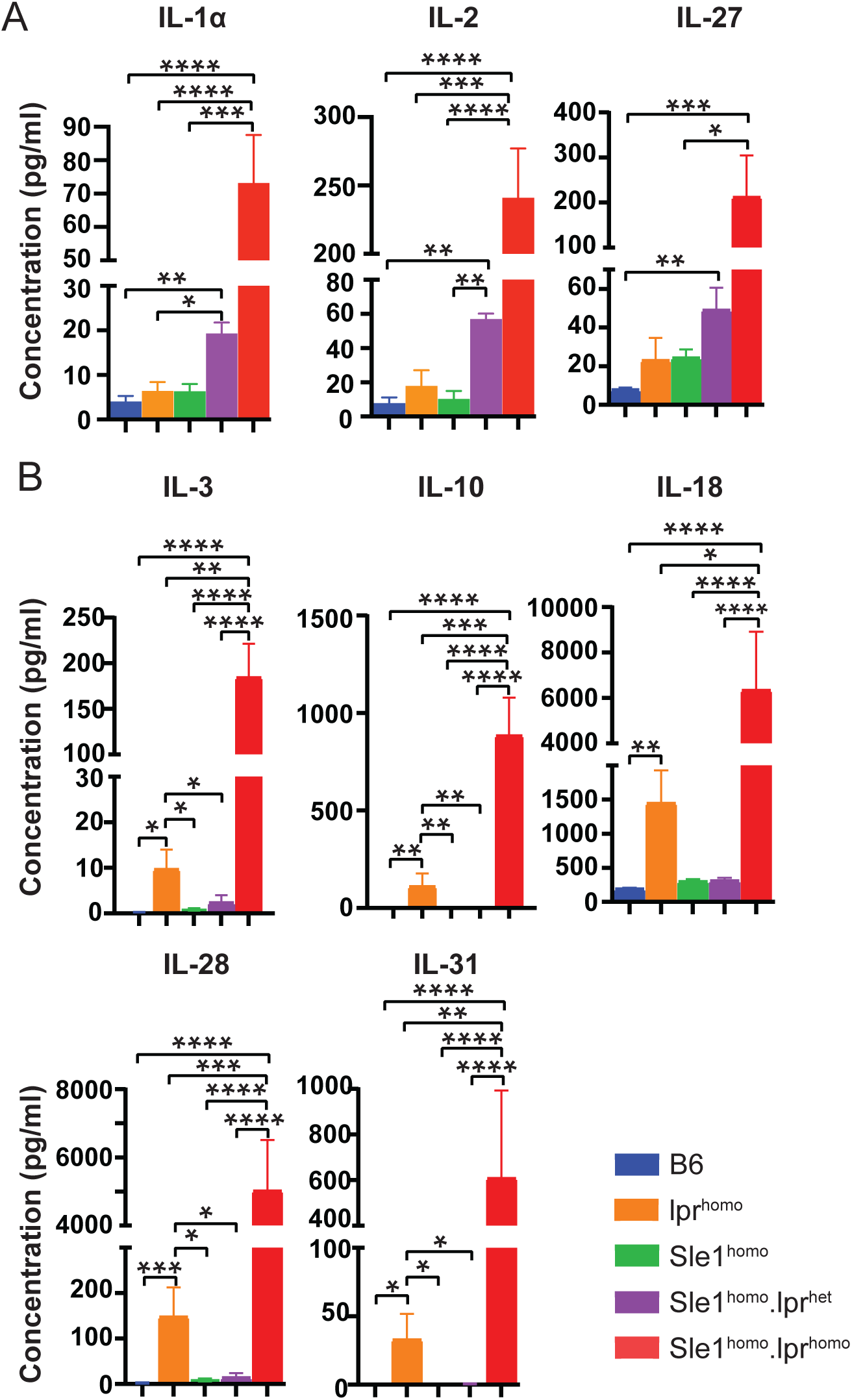
Impact of *FAS^lpr^*zygosity on serum cytokine levels. Serum levels of cytokine/chemokines in B6 and B6 congenic mice at 2-3 months of age were measured by Luminex assays. (A) serum cytokines levels of IL-1*a,* IL-2, and IL-27 showing phenotype driven by *FAS^lpr^* zygosity, in concert with *sle1* homozygosity. (B) serum levels of cytokines showing phenotype driven by homozygosity of *FAS^lpr^* zygosity. Tested B6 congenic mice include lpr^homo^, Sle1^homo^, Sle1^homo^.lpr^het^, and Sle1^homo^.lpr^homo^. N=5-6/strain. *p<0.05, **p<0.01, ***p<0.001 and ****p<0.0001.

### 3.5 Skewed Th1 differentiation in Sle1^homo^.lpr^het^ mice

Among the limited cytokines elevated in sle1^homo^.lpr^het^ mice, IL-2 and IL-27 have been reported to be indicators of Th1-type immune response [20, 21]. Thus, we hypothesized that heterozygosity at *Fas^lpr^* on the B6.sle1^homo^ background may alter CD4^+^ helper T cell differentiation, which may contribute to autoimmune responses without the profound effect on lymphoproliferation. To test this hypothesis, we isolated naïve CD4^+^ T cells from B6 and sle1^homo^.lpr^het^ mice, stimulated with anti-CD3 and anti-CD28 antibodies, then cultured T cells in different types of medium to polarize towards Th1, Th2, and Th17 differentiation. The medium containing only IL-2 (Th0) was used to support T cell growth without polarization. Phenotypes of differentiated CD4^+^ T cells were determined by the production of IFN-γ, IL-4, and IL-17, key effector cytokines for Th1, Th2, and Th17 cells, respectively. As shown in Figure 6A, an increased percentage of IFN-γ^+^ cells in T cells under Th1 and Th17 polarization conditions was observed in sle1^homo^.lpr^het^ mice (Fig 6A). There were no significant changes in IL-4^+^ cells under the Th2 polarization condition between these two strains (Fig 6A). Moderately decreased IL-17^+^ cells under Th17 differentiation were also noted in sle1^homo^.lpr^het^ mice (Fig 6A). Consistent with the flow cytometry-based analysis, ELISA results demonstrated that sle1^homo^.lpr^het^ CD4^+^ T cells in the presence of Th1-, Th17-polarization medium or IL-2 only medium produced more IFN-γ than counterpart controls from B6 mice (Fig 6B). As expected, increased IL-4 and IL-17 production was observed in T cells differentiated in the Th2-polarization medium and Th17-polarization medium, respectively. However, IL-4 production by Th2-differentiated sle1^homo^.lpr^het^ T cells and IL-17 production by Th17-differentiated sle1^homo^.lpr^het^ T cells were significantly lower than those from B6 controls (Fig 6C). Together, our data suggest that partial loss of *FAS* function on B6.sle1^homo^ background preferentially drives differentiation of naïve CD4^+^ T cells towards Th1 cells upon TCR stimulation.

**Figure 6.**
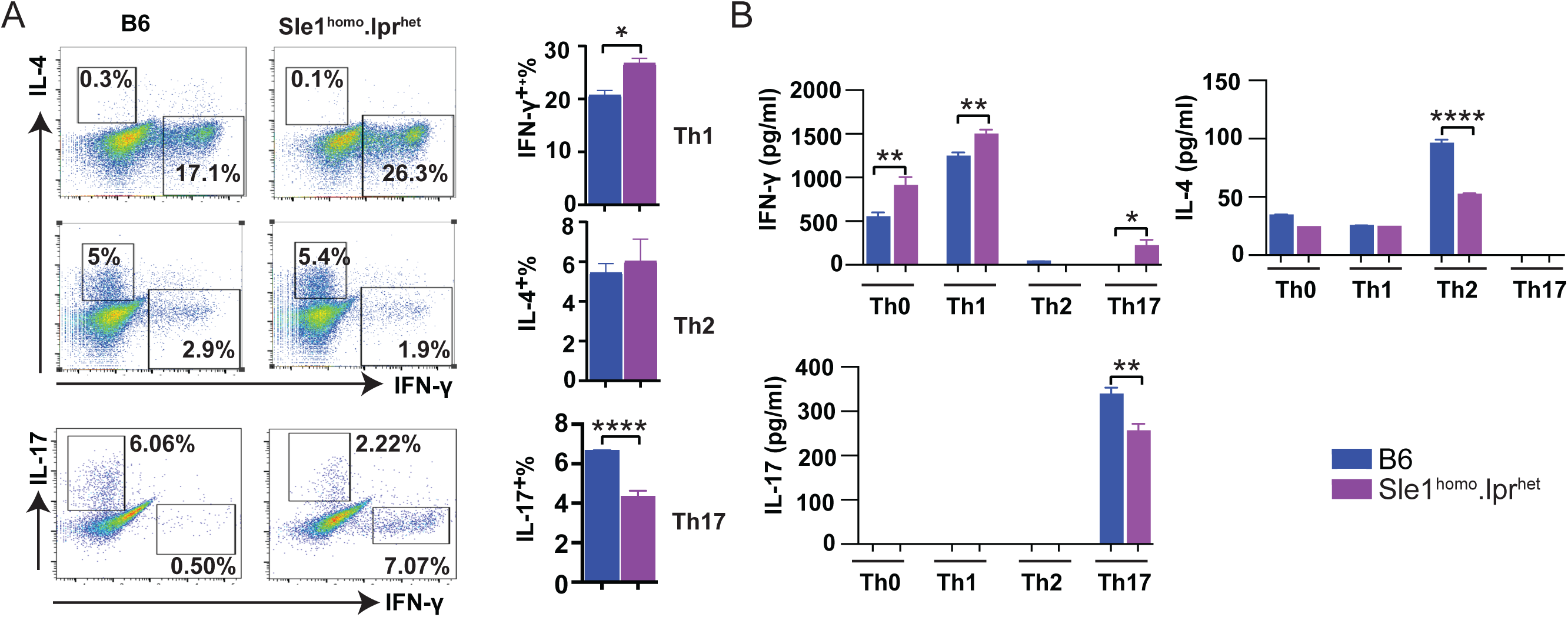
T cell differentiation changes in Sle1^homo^.lpr^het^ mice. Naïve CD4^+^ T cells were isolated from spleens of B6 and Sle1^homo^.lpr^het^ mice and were stimulated with 2.5 μg/ml of anti-CD3e and 1 μg/ml of anti-CD28 with IL-2 only or under culture conditions for Th1, Th2, and Th17 differentiation. Cultured T cells were restimulated and used to evaluate IFN-γ, IL-4, and IL-17 production by flow cytometry analysis and ELISA. (A) Representative flow cytometry results and bar plots showing the percentage of IFN-γ^+^, IL-4^+^, and IL-17^+^ T cells in different culture conditions (B) ELISA data showing the concentration of IFN-γ, IL-4 and IL-17 in culture supernatant in restimulated T cells from different culture conditions. N=3/strain; Mice from both genders were included. *p<0.05, **p<0.01, ***p<0.001 and ****p<0.0001.

## 4. Discussion

A broad range of immunological dysfunctions involving both innate and adaptive immune systems have been identified in SLE patients [2], including abnormal serum levels of cytokines [22], altered complement activation [23], production of pathogenic autoantibodies [24] and hyperactive of T and B cells[25], as recently reviewed [26]. Although many murine models have been generated to study the molecular and cellular mechanism of lupus [5], there is no single experimental model that mirrors all phenotypes present in human SLE. Introducing lupus susceptibility loci on the B6 background has resulted in establishing a series of B6 congenic lupus strains [4, 7, 10, 27-31]with distinct phenotypes. While B6 monocongenic lupus strains exhibit limited or no autoimmune symptoms or renal diseases, introgression of multiple lupus susceptibility loci leads to massive production of autoantibodies and fatal lupus nephritis as previously reported in the B6.*sle1.sle2.sle3* and B6.sle1^homo^.lpr^homo^ strains [10, 29]. However, these two strains display severe hyperproliferation of T and B cells. Therefore, phenotype characterization of mice bearing one or multiple lupus susceptibility loci, particularly in B6 congenic background, can not only identify the link between genetic alterations and immunological dysfunctions but also provide a series of congenic lupus models to better represent the kaleidoscope of clinical manifestations in human SLE.

Whereas *Fas^lpr^* impairs cell apoptosis and activation-induced cell death, *sle1* breaches B cell tolerance, leading to the emergence of a chromatin-targeted autoantibody repertoire, insufficient to elicit significant renal damage [32-34]. Here, we test whether impairing lymphocyte apoptosis (using the *Fas^lpr^* allele) can transition the humoral autoimmunity seen in *sle1* mice into more severe lupus-like disease states. As anticipated, sle1^homo^.lpr^het^ mice starting at 4-mo age exhibited elevated ANA, proteinuria, and mild glomerulonephritis. There was no evidence of splenomegaly or lymphadenopathy in this strain. Unlike in sle1^homo^.lpr^homo^ mice, percentages of proliferative T and B cells in sle1^homo^.lpr^het^ mice were comparable with B6 controls.

Interestingly, immune profiling of immune cells from sle1^homo^.lpr^het^ mice revealed the effects of *Fas^lpr^* in upregulating serum levels of IL-2 and IL-27, two Th1-associated cytokines, resulting in Th1 skewed T cell differentiation. These results suggest that the heterozygous absence of *FAS* interacting with *sle1* can augment anti-chromatin autoimmunity by upregulating Th1-based immune responses.

Early studies in mouse models showed that autoimmune responses in murine lupus models can be mitigated by T cell depletion [35, 36]. T cell abnormalities are also frequently observed in SLE patients [37, 38]. While CD8^+^ T cells have been suggested to directly contribute to autoimmune-mediated tissue damage [39, 40], data also reveal that pathogenic CD4^+^ T cells promote chronic inflammatory responses through releasing cytokines and elevate autoantibody levels through helping self-reactive B cells in lupus [41]. Here, our studies showed immune responses mediated by CD4^+^ T cells from sle1^homo^.lpr^het^ mice are skewed to Th1-type, and increased levels of cytokines associated with Th1 responses are also noted in this strain. Even under appropriate cytokine environments, sle1^homo^.lpr^het^ naïve CD4^+^ T cells exhibited significantly reduced Th2 and Th17 differentiation. Results from this new lupus strain are consistent with clinical data indicating increased IFN-γ production and lowered IL-4 production in SLE [42]. Therefore, our data not only provide direct evidence to link pathogenic Th1 responses with lupus-like autoimmunity but also support the usage of this strain to study Th1-related mechanistic studies or therapeutic approaches. In the same vein, we also observed the gene dose effect of *FAS* on elevating serum IL-27 levels. IL-27, produced mainly by dendritic cells and macrophages, belongs to the IL-12 family [43]. Upon Toll-like Receptor (TLR) stimulation, IL-27 has been reported to induce IFN-γ production via the Jak-Stat pathway and serve as an early initiator of Th1 responses [21, 43]. Development of arthritis was delayed in IL-27R knockout (KO) mice, and IL-27R KO only alters the frequency of IFN-γ^+^ T cells, not IL-4^+^ or IL-17^+^ T cells [44]. Recently, the hyperlipidemia-TLR4-IL27-T_FH_ cell axis was implicated in promoting atherosclerosis in SLE patients [45], and certain IL-27 gene polymorphisms are noted to be associated with SLE [46].

Perhaps most importantly, the newly generated strain lacks the profound lymphoproliferation described in the traditionally studied sle1^homo^.lpr^homo^ mice, which are also regarded as a mouse model of chronic lymphocytic leukemia. Since the new strain does not die prematurely due to lymphosplenomegaly, this allows one to age these mice further till autoimmunity and renal disease gradually set in, reminiscent of patients with lupus. Since the sle1^homo^.lpr^het^ strain bears only one allele of the mutant *FAS* allele and is on the C57BL/6 genetic background, this strain is ideal for investigating the additive effect of other immune genes already backcrossed onto the same genetic background. Hence, the newly generated sle1^homo^.lpr^het^ strain constitutes a novel mouse model of lupus, ideal for genetic studies, autoantibody repertoire investigation, and Th1 effector cell skewing.

## Abbreviations

SLE: Systemic Lupus Erythematous
B6: C57BL/6
lpr^homo^: B6.*Fas^lpr/lpr^*
sle1^homo^: B6.*Sle1/Sle1*
sle1^homo^.lpr^het^: B6.*Sle1/Sle1.Fas^lpr^*^/+^
sle1^homo^.lpr^homo^: B6.*Sle1/Sle1.Fas^lpr/lpr^*
IFN: interferon
H&E: Hematoxylin and Eosin
Anti-dsDNA: Anti-double-stranded DNA
ANA: anti-nuclear antibody
BUN: Blood Urea Nitrogen
FFPE: Formalin-Fixed and Paraffin-Embedded
mo: months
hr: hour
GN: glomerulonephritis
KO: Knockout
Th: T helper
TLR: Toll-like Receptor
OD: optical density
NS: no statistical significance
NA: Not available

## Authors’ Contributions

**Conception and Design**: C. Mohan and W. Peng

**Acquisition of data (provided required animals, cells, animal samples, etc.):** R. Bohat, X. Liang, C. Xu, Y. Chen, N. Zheng, A. Guerrero, R. Jaffery, N. Egan, A. Robles, Y. Du, C. Mohan, and W. Peng

**Analysis and interpretation of data (statistical analysis):** R. Bohat, X. Liang, Y. Chen, M. Hicks, X. Chen, C. Mohan, and W. Peng

**Writing and/or revision of the manuscript:** R. Bohat, C. Mohan, and W. Peng

**Study supervision**: W. Peng

