## Supplementary Figures and Tables for "*FAS^lpr^* gene dosage tunes the extent of lymphoproliferation and T cell differentiation in lupus"

Bohat R, *et al.* Supplementary Figure 1

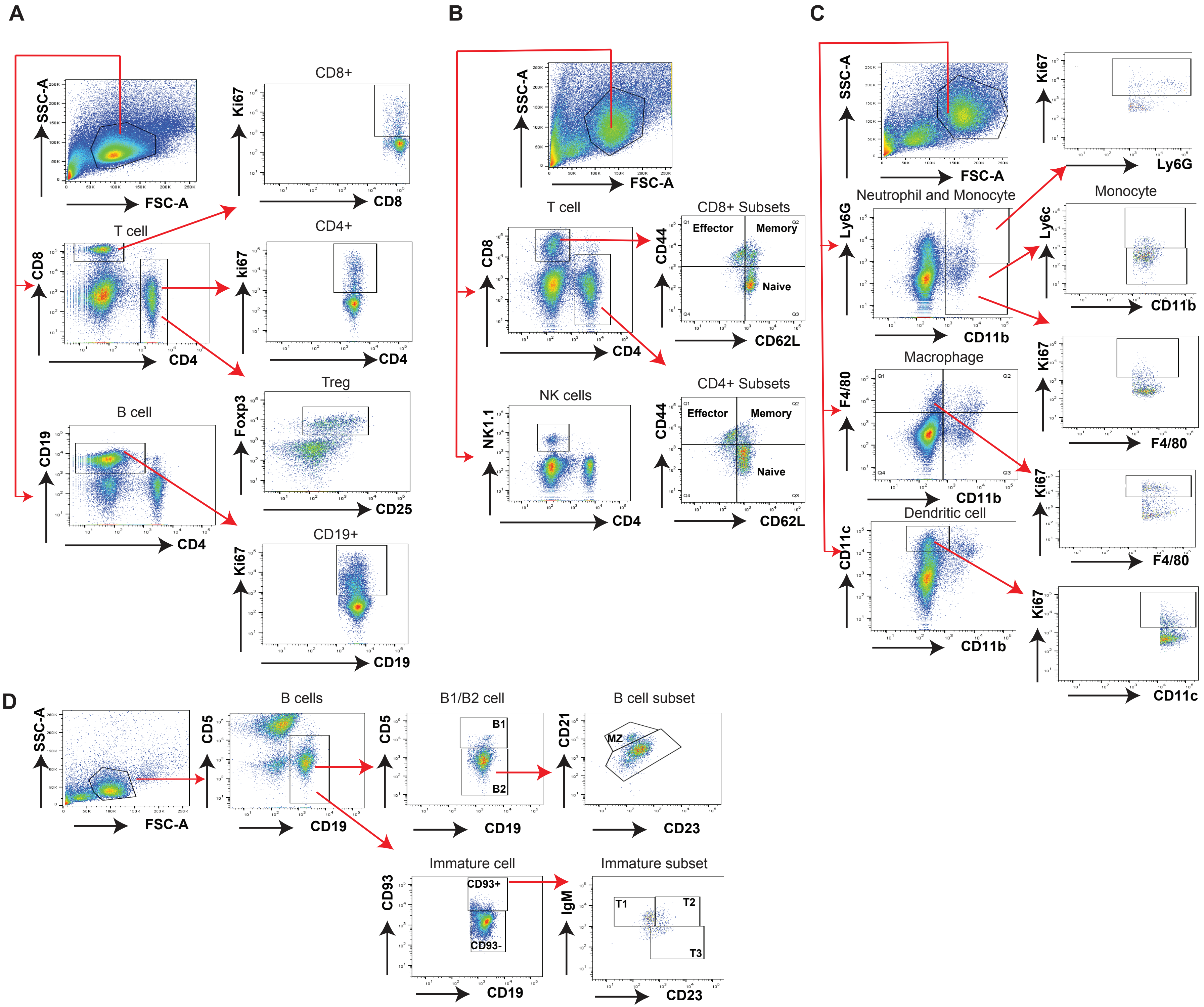

**Bohat R, *et al.* Supplementary Figure 2**

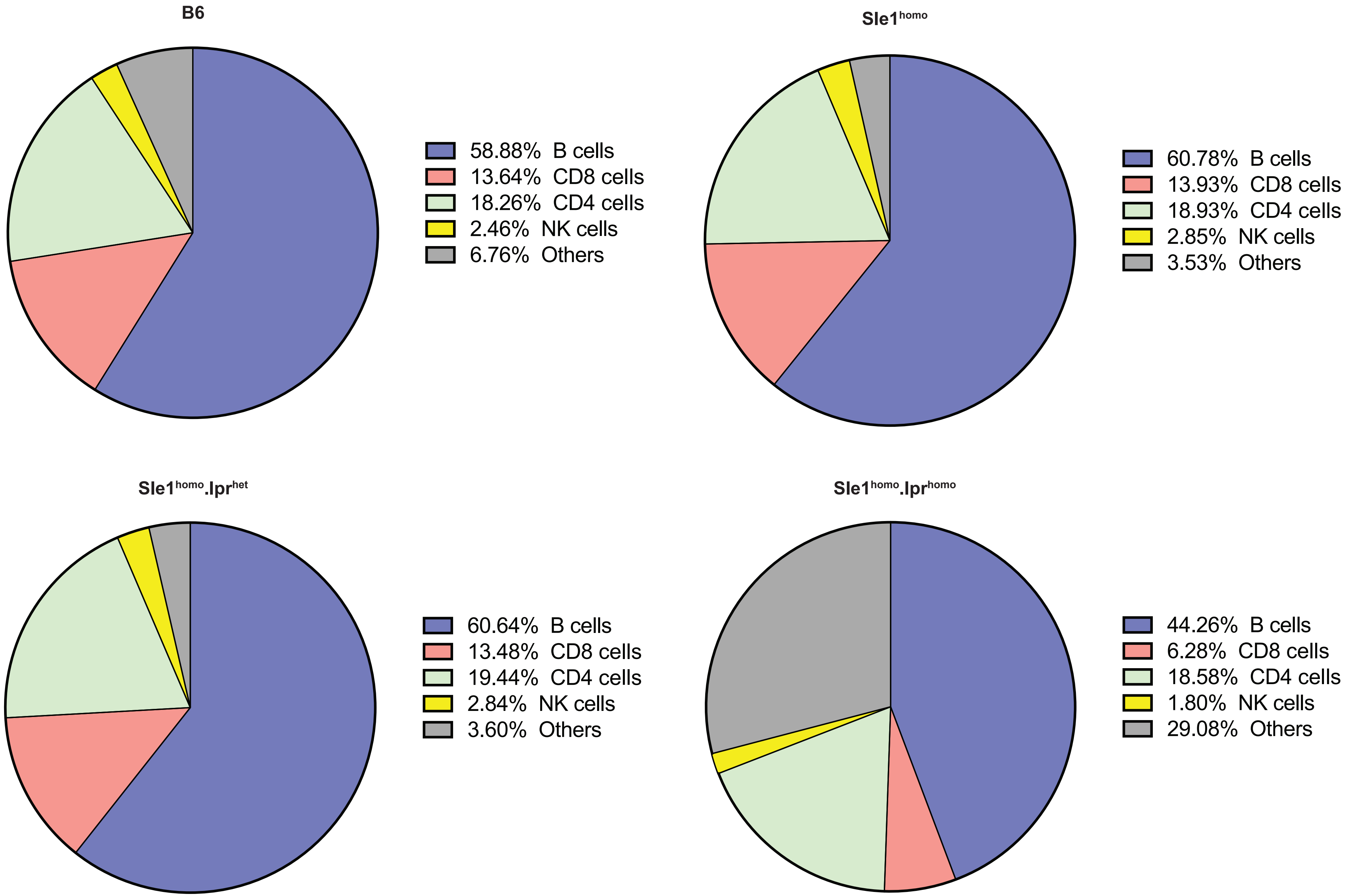

**Bohat R, *et al.* Supplementary Figure 3**

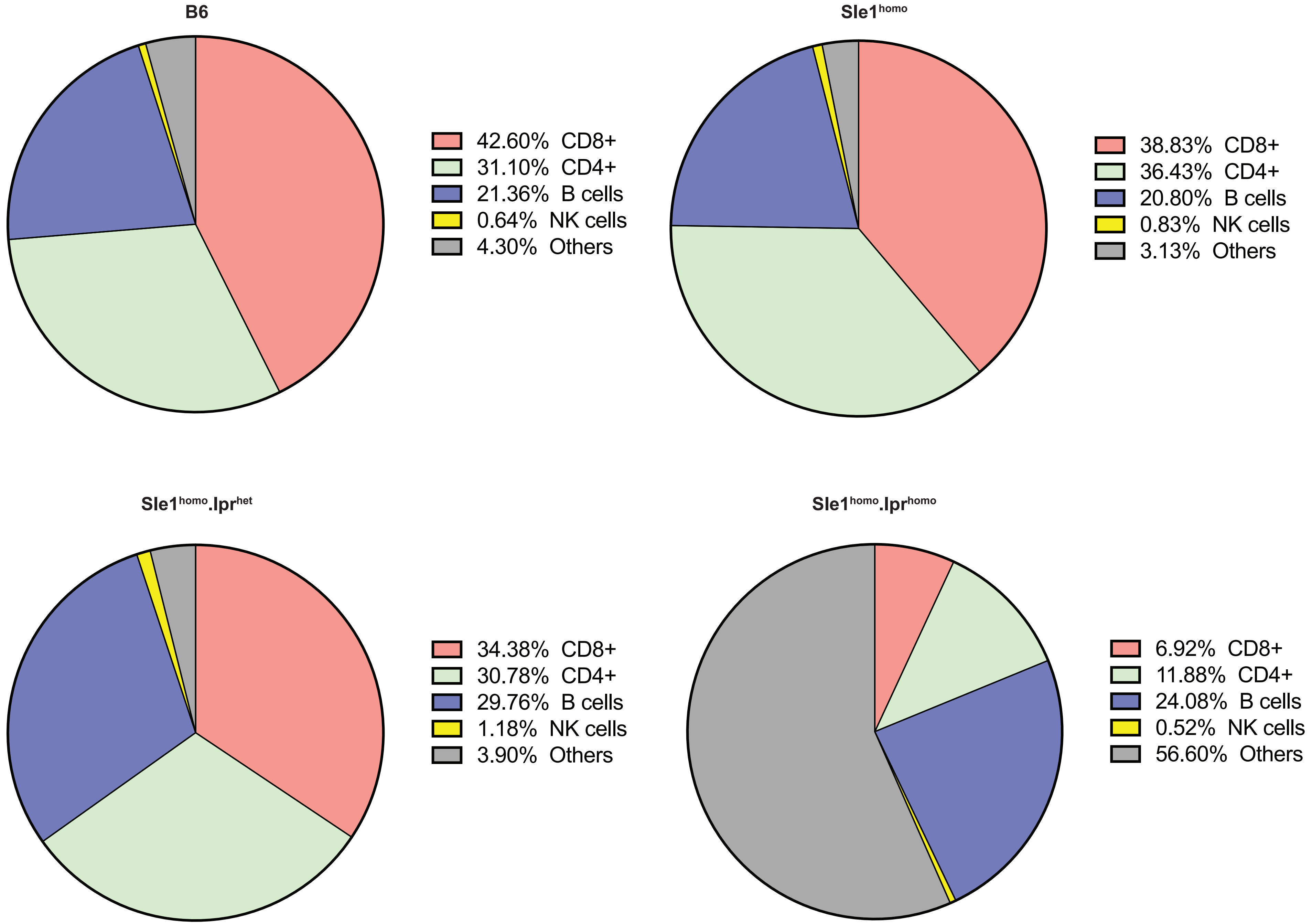

| **Supplementary Table 1** | | | |
|---|---|---|---|
| **anti-dsDNA OD (Mean±SEM)** | | | |
| | **Strain** | **Female** | **Male** |
| **2-3 Months** | **B6** | 0.07±0.003 | 0.07±0.003 |
| | **$Sle1^{homo}$** | 0.08±0.01 | 0.08±0.002 |
| | **$Sle1^{homo}lpr^{het}$** | 0.08±0.003 | 0.08±0.004 |
| | **$Sle1^{homo}lpr^{homo}$** | 0.2±0.07 | 0.22±0.03 |
| **4-8 Months** | **B6** | 0.09±0.007 | 0.09±0.02 |
| | **$Sle1^{homo}$** | 0.12±0.02 | 0.08±0.002 |
| | **$Sle1^{homo}lpr^{het}$** | 0.2±0.04 | 0.1±0.006 |
| | **$Sle1^{homo}lpr^{homo}$** | NA | NA |
| **10-12 Months** | **B6** | 0.11±0.01 | 0.08±0.006 |
| | **$Sle1^{homo}$** | 0.19±0.08 | 0.18±0.07 |
| | **$Sle1^{homo}lpr^{het}$** | 0.23±0.004 | 0.11±0.03 |
| | **$Sle1^{homo}lpr^{homo}$** | NA | NA |

| Supplementary Table 2 | | | |
|---|---|---|---|
| **Total protein in mg (Mean±SEM)** | | | |
| | **Strain** | **Female** | **Male** |
| **2-3 Months** | **B6** | 69±26.2 | 276.6±51.7 |
| | **$Sle1^{homo}$** | 204.9±56.3 | 377.2±72 |
| | **$Sle1^{homo}lpr^{het}$** | 179±37.5 | 370±112.7 |
| | **$Sle1^{homo}lpr^{homo}$** | 333.14±88.07 | 671.3±338.1 |
| **4-8 Months** | **B6** | 163.6±14.6 | 170.7±67.7 |
| | **$Sle1^{homo}$** | 144.2±61.7 | 175.1±39.3 |
| | **$Sle1^{homo}lpr^{het}$** | 128.3±11.4 | 611.5±111.3 |
| | **$Sle1^{homo}lpr^{homo}$** | NA | NA |
| **10-12 Months** | **B6** | 166.2±17.5 | 169±70 |
| | **$Sle1^{homo}$** | 113.3±33.2 | 573.4±NA |
| | **$Sle1^{homo}lpr^{het}$** | 174.1±6.0 | 323±60 |
| | **$Sle1^{homo}lpr^{homo}$** | NA | NA |

| Supplementary Table 3 | | | |
|---|---|---|---|
| BUN in mg/dL (Mean±SEM) | | | |
| | **Strain** | **Female** | **Male** |
| **2-3 Months** | **B6** | 6.2±1.6 | 12.8±2.6 |
| | **Sle1$^{homo}$** | 15.2±2.4 | 11.4±0.32 |
| | **Sle1$^{homo}$lpr$^{het}$** | 22±7 | 18.2±1.2 |
| | **Sle1$^{homo}$lpr$^{homo}$** | 27.8±11.6 | 16±1 |
| **4-8 Months** | **B6** | 7.6±1.2 | 12.8±3.2 |
| | **Sle1$^{homo}$** | 14.2±1.4 | 12.9±1.2 |
| | **Sle1$^{homo}$lpr$^{het}$** | 19.3±4 | 13.3±1.2 |
| | **Sle1$^{homo}$lpr$^{homo}$** | NA | NA |
| **10-12 Months** | **B6** | 10.0±1.05 | 12.0±0.9 |
| | **Sle1$^{homo}$** | 18.2±1.4 | 15.9±0.5 |
| | **Sle1$^{homo}$lpr$^{het}$** | 16.5±0.5 | 15.5±1.4 |
| | **Sle1$^{homo}$lpr$^{homo}$** | NA | NA |

**Supplementary Table 4**

| Absolute number (Mean±SEM) | | | | | p value B6 vs. | | | p value Sle1$^{homo}$lpr$^{het}$ Vs. |
|---|---|---|---|---|---|---|---|---|
| **Cell type** | **B6** | **Sle1$^{homo}$** | **Sle1$^{homo}$lpr$^{het}$** | **Sle1$^{homo}$lpr$^{homo}$** | **Sle1$^{homo}$** | **Sle1$^{homo}$lpr$^{het}$** | **Sle1$^{homo}$lpr$^{homo}$** | **Sle1$^{homo}$lpr$^{homo}$** |
| **CD8** | (5.6±0.3)x10$^{6}$ | (9.6±1.3)x10$^{6}$ | (10.2±1.0)x10$^{6}$ | (6.8±1.5)x10$^{6}$ | NS | * | NS | NS |
| **CD4** | (7.6±0.7)x10$^{6}$ | (13.0±1.6)x10$^{6}$ | (14.8±1.5)x10$^{6}$ | (19.6±3.2)x10$^{6}$ | NS | NS | ** | NS |
| **B cells** | (24.2±1.4)x10$^{6}$ | (42.6±6.8)x10$^{6}$ | (45.6±3.7)x10$^{6}$ | (50.8±14.6)x10$^{6}$ | NS | NS | NS | NS |
| **NK cells** | (7.9±0.5)x10$^{4}$ | (18.8±4.1)x10$^{4}$ | (25.3±2.7)x10$^{4}$ | (51.9±4.2)x10$^{4}$ | NS | ** | **** | **** |
| **Monocytes** | (4.9±0.4)x10$^{4}$ | (15±0.4)x10$^{4}$ | (15.7±4.1)x10$^{4}$ | (33.5±6.0)x10$^{4}$ | NS | NS | ** | * |
| **Neutrophil** | (11.1±0.8)x10$^{4}$ | (9.3±0.6)x10$^{4}$ | (24.8±6.0)x10$^{4}$ | (31.3±6.2)x10$^{4}$ | NS | NS | NS | NS |
| **Macrophage** | (13.5±5.9)x10$^{4}$ | (10.7±3.2)x10$^{4}$ | (33.2±8.3)x10$^{4}$ | (66.5±7.6)x10$^{4}$ | NS | NS | ** | * |
| **DC** | (8.1±0.6)x10$^{4}$ | (15.5±2.0)x10$^{4}$ | (20.2±0.6)x10$^{4}$ | (134.8±24.5)x10$^{4}$ | NS | NS | *** | *** |

**Supplementary Table 5**

| Absolute number (Mean±SEM) | | | | | p value B6 vs. | | | p value Sle1$^{homo}$lpr$^{het}$ Vs. |
|---|---|---|---|---|---|---|---|---|
| **Cell type** | **B6** | **Sle1$^{homo}$** | **Sle1$^{homo}$lpr$^{het}$** | **Sle1$^{homo}$lpr$^{homo}$** | **Sle1$^{homo}$** | **Sle1$^{homo}$lpr$^{het}$** | **Sle1$^{homo}$lpr$^{homo}$** | **Sle1$^{homo}$lpr$^{homo}$** |
| **CD8** | (6.6±0.9)x$10^5$ | (8.0±1.7)x$10^5$ | (6.4±2.7)x$10^5$ | (14.4±3.2)x$10^5$ | NS | NS | NS | NS |
| **CD4** | (4.9±0.8)x$10^5$ | (7.3±1.3)x$10^5$ | (6.1±2.8)x$10^5$ | (26.4±8.2)x$10^5$ | NS | NS | * | * |
| **B cells** | (3.3±0.5)x$10^5$ | (4.5±1.3)x$10^5$ | (6.1±3.0)x$10^5$ | (44.5±11.6)x$10^5$ | NS | NS | *** | ** |
| **NK cells** | (6.5±0.5)x$10^2$ | (11.5±1.2)x$10^2$ | (14.9±4.6)x$10^2$ | (105.6±15.8)x$10^2$ | NS | NS | **** | **** |
| **Monocytes** | (5.5±0.3)x$10^3$ | (4.4±1.2)x$10^3$ | (10.4±3.7)x$10^3$ | (14.3±4.8)x$10^3$ | NS | NS | NS | NS |
| **Neutrophil** | (2.2±0.1)x$10^3$ | (1.8±0.5)x$10^3$ | (6.4±2.5)x$10^3$ | (27.9±7.5)x$10^3$ | NS | NS | * | * |
| **Macrophage** | (4.7±0.9)x$10^2$ | (2.3±0.4)x$10^2$ | (12.4±5.3)x$10^2$ | (149.9±51.2)x$10^2$ | NS | NS | * | * |
| **DC** | (1.6±0.2)x$10^4$ | (1.1±0.1)x$10^4$ | (3.1±1.0)x$10^4$ | (83.6±22.1)x$10^4$ | NS | NS | ** | ** |

**Legends: Supplementary Figures and Tables**

**Supplementary Figure 1. Gating strategies of flow cytometry analysis for immune profiling.** (A) The gating strategy to determine the percentage and proliferation of T and B cells. (B) the gating strategy to determine percentages of effector/memory/naïve T cells and NK cells. (C) the gating strategy to determine the percentage and proliferation of macrophage and myeloid cells. (D) the gating strategy used to determine B cell subsets.

**Supplementary Figure 2. Pie chart showing the distribution of major immune subsets within the spleen.** Averages of each immune subset in spleens from B6 and B6 congenic mice were used to plot the pie chart.

**Supplementary Figure 3. Pie chart showing the distribution of major immune subsets within the lymph nodes.** Averages of each immune subset in lymph nodes from B6 and B6 congenic mice were used to plot the pie chart.

**Supplementary Table 1. Serum levels of IgG autoantibodies to dsDNA in B6 and B6 congenic mice.** Blood samples were collected from both female and male mice at three different ages: 2-3 months (mo), 4-8 mo, and 10-12 mo. The average of anti-dsDNA serum levels in male and female mice are listed. OD: optical density; NA: Not available.

**Supplementary Table 2. 24-hr total urine protein amount in B6 and B6 congenic mice.** The average of 24-hr urine protein levels in male and female mice are listed. NA: Not available.

**Supplementary Table 3. Blood Urea Nitrogen (BUN) level in B6 and B6 congenic mice**. The average of BUN levels in male and female mice are listed. NA: Not available.

**Supplementary Table 4. Absolute numbers of immune cells in spleen from B6 and B6 congenic mice.** Averages of the numbers of immune cells in each strain and the status of statistical significance between

B6 and B6 congenic mice were listed. NS: no statistical significance. *p<0.05, **p<0.01, ***p<0.001 and ****p<0.0001.

**Supplementary Table 5. Absolute number of immune cells in lymph nodes from B6 and B6 congenic mice.** Averages of numbers of immune cells in each strain and the status of statistical significance between B6 and B6 congenic mice were listed. NS: no statistical significance. *p<0.05, **p<0.01, ***p<0.001 and ****p<0.0001.
